# When the heart meets the mind Exploring the brain-heart interaction during time perception

**DOI:** 10.1101/2023.09.20.558404

**Authors:** Shiva Khoshnoud, David Leitritz, Meltem Çinar Bozdağ, Federico Alvarez Igarzábal, Valdas Noreika, Marc Wittmann

## Abstract

It has been hypothesized that time estimation relies on bodily rhythms and interoceptive signals, such as heartbeats. Extending previous research demonstrating this connection, we provided further electrophysiological evidence that the brain registers heartbeats while tracking time intervals. We evaluated the heartbeat-evoked potential (HEP) and the contingent negative variation (CNV) during an auditory duration-reproduction task with intervals lasting 4, 8, and 12 seconds and a control reaction-time task employing the exact durations. The (interoceptive) Self-Awareness Questionnaire (SAQ) and the heartbeat-counting task (HCT) were administered. The SAQ, but not the HCT scores, correlated with the mean reproduced durations for 4s, 8s, and 12s intervals: the higher the SAQ score (a stronger awareness of bodily signals), the longer the duration reproductions and the more accurate the timing behavior. The HEP amplitude within 130-270 ms (HEP1) and 470-520 ms (HEP2) after the R peak was smaller for the 4s interval than for the 8s and 12s intervals. This is a timing-specific effect, as the HEP amplitude did not differ for intervals in the reaction-time task. A ramp-like increase in HEP2 amplitudes was significant for the duration-encoding phase of the timing task, but not for the control reaction-time task. The HEP2 increase within the reproduction phase of the timing task correlated significantly with the reproduced durations for the 8s and the 12s intervals. The larger the registered increase in the HEP2, the greater the under-reproduction of the estimated duration. The initial and late CNV components were significantly more negative during the encoding phase than during the reaction-time task. Given the link between these components with attention modulation and temporal memory, we interpret the CNV findings as indicating greater executive resources oriented towards time. We conclude that interoceptive awareness (SAQ) and state-like cortical responses to the heartbeat (HEP) predict duration reproductions, emphasizing the notion of the embodiment of time.

## Introduction

How and where event duration is processed in the brain remains an unsolved question. Timing mechanisms in the brain are essential for an organism to represent temporal regularities in the environment and the temporal metrics of events [1]. There exists a strong consensus on basic neural processes regarding space perception and spatial navigation related to entorhinal and hippocampal structures [2], a work for which the Nobel Prize for Physiology was awarded in 2014. The underlying neural structures pertaining to the sense of time await such a consensus.

Many neuroimaging studies have attempted to delineate the neural basis of the human sense of time. Meta-analyses of functional magnetic resonance imaging (fMRI) studies implicate a relatively stronger involvement of subcortical processes with stimuli in the sub-second duration range; relatively stronger activation of cortical regions is registered with supra-second durations [3]. One meta-analysis based on the activation-likelihood estimation of fMRI experiments identified a network of brain areas involved in time perception which encompassed the supplementary motor area (SMA), the intraparietal sulcus, the inferior frontal gyrus, the insular cortex, and the basal ganglia [1]. The multitude of neural activation sites identified suggests that there is not just one mechanism in the brain responsible for processing duration (a master clock) but that there could be distributed cerebral networks [4]. Specific dynamic properties of a given neural circuit could possibly govern the processing of duration [5]. Different functional subcomponents for tracking time could explain these distributed activations [6,7]. Each identified parallel brain circuit might predominantly be activated when required by different tasks, modalities, and the duration processed [4].

Teghil et al. [8] indicate how parallel timing systems can be activated according to the task. In their experimental setup, temporal intervals of several seconds duration were filled with regularly or irregularly presented sequential visual stimuli. This setup was designed to distinguish between two conditions with regular external cues (which would help process duration) and no external cues (irregular condition). Connectivity of the right posterior insula with other brain regions was modulated by the participants’ degrees of interoceptive awareness (assessed with the Self-Awareness Questionnaire (SAQ; Longarzo et al., 2015), which only correlated positively with performance in the irregular condition. The interpretation favored by the authors was that the ability to sense their own body states enabled participants to be more accurate in the duration-reproduction task when no regular external cues were available. The two most recent meta-analyses of more than 100 neuroimaging experiments using activation-likelihood estimates for brain structures converge on only two sites: the (pre-) SMA and bilateral insula [10,11]. These results correlate with the idea that subjective time is driven by sensorimotor or enactive (SMA) processing and interoceptive or embodied (insula) experience [11].

The notion of a link between bodily states and subjective time as being related to insular-cortex function was proposed on conceptual grounds over a decade ago [12]. Accumulating empirical evidence had provided increasing support for this hypothesis [13]. A conceptual connection between body awareness and subjective time was established because the two have access to a common neural system, the interoceptive system, which includes the insular cortex. Utilizing the duration-reproduction paradigm with multiple-second tone intervals, which was employed in the present study, a ramp-like increase in fMRI activation terminated with the end of the auditory stimulus during the presentation of the first interval, the encoding phase (climbing activity in the dorsal posterior insula), and the second interval, when the duration of the first stimulus had to be reproduced (climbing activity in the anterior insula) [14,15]. Complementing the interpretation of Teghil and colleagues [8,16], we concur that since no regularly occurring external cues were present in the specific duration-reproduction task, and a secondary working-memory task controlled chronometric counting, participants had to largely rely on body-related processes to be able to time their behavior. To test the hypothesis of the involvement of the autonomic nervous system in time perception, peripheral-physiological measures were employed in a further study while participants performed the duration-reproduction task. Cardiac periods increased, and skin-conductance levels decreased steadily in the range of several seconds in the auditory [17] and visual modalities [18] during the encoding and the reproduction phases of the timing task. Similarly, duration timing in the millisecond time range has been shown to be influenced by the participants’ beat-by-beat cardiac dynamics [19,20].

These findings suggest that interoceptive processes underlie time perception in the range of milliseconds to seconds. In contextualizing these and the following empirical findings, it is likely that interoceptive signals are more strongly required to judge time duration when no external cues are present. It has been shown that the phase locking between the cardiac cycle and the start/stop interval signals in the duration-reproduction task enhances the accuracy of duration reproduction [21]. The heartbeat-evoked potential (HEP) recorded in the brain is a measure of interoceptive cortical processing, which covaries with bodily self-consciousness [22,23]. The heartbeat-perception accuracy in the heartbeat-counting task (HCT) measures interoceptive sensitivity (see below). The HEP amplitude and the HCT accuracy have been shown to predict time estimates for intervals lasting 120 seconds [24]. However, in a different study, when the duration of presented film clips with complex and dynamic contents had to be judged, the perceptual visual content and not the physiological signals, such as the heart rate, were predictive of the duration estimates [25].

One measure of interoceptive sensitivity is the heartbeat-counting task or HCT, as initially conceptualized by Schandry [26]. Individuals are asked to count their heartbeats without monitoring their pulse and report the number at the end of a defined interval. A more accurate heartbeat report can be interpreted as stemming from better body awareness. In accordance with the body-state hypothesis of time perception, in one study, those participants who performed more accurately in the duration-reproduction task (with less deviation from objective duration) also did so in the heartbeat-counting task [17]. This finding, however, was not replicated in a follow-up study with slightly older (middle-aged) participants; the heartbeat-counting scores were not related to subsequent timing performance in a duration-reproduction task [18]. Methodological criticism related to the heartbeat-counting task must be discussed here. It has been argued that participants could guess their approximate heart rates and thus perform the task quite accurately, even without sensing their heartbeats [27]. Therefore, the need for other objective psychophysiological tasks when assessing interoceptive accuracy has been raised [28]. Nevertheless, the two positive findings of a relationship between heartbeat-counting accuracy and time estimation [17,24] encouraged us to utilize this measure of interoception in combination with an auditory duration-reproduction task.

Previous fMRI and psychophysiological studies showed climbing activity across the intervals to be timed in human participants [13]. Animal timing research has similarly shown that climbing neural activity in several brain regions is associated with timed motor behavior [29–31]. Human electroencephalography (EEG) studies have also demonstrated how climbing neural activation in the SMA accompanies event timing [32]. While participants estimate the duration of presented intervals, a slow negative electrophysiological shift, referred to as the contingent negative variation (CNV), has been recorded. The CNV appears in explicit (duration estimation) and implicit (reaction time) timing tasks [33]. The CNV has been suggested to represent an accumulation mechanism in duration timing as proposed in cognitive pacemaker–accumulator models [32,34]. Theoretical interval-timing models suggest that climbing neural activity results from neural firing-rate adaptations in individual neurons and neuronal populations [35–37]. The idea that the CNV refers to an explicit timing mechanism (an internal clock) has been challenged by conceptual deliberations; it may instead reflect general processes of expectation, temporal memory, and response preparation [38–40].

We combined different measures identified to correlate with time perception in the present study. We used the auditory duration-reproduction task, where participants are requested to reproduce the duration of presented tones. The secondary working-memory task prevented participants from counting. A reaction-time task was used as a control. We recorded EEGs and registered the heartbeats while participants performed the task to analyze the CNV and the HEP components. Interoceptive performance was evaluated with the participants’ performance in the classic heartbeat-counting task and with the assessment of the trait-like interoceptive awareness questionnaire (SAQ), which has been shown to be sensitive to performance in the duration-reproduction task [8]. We hypothesized that a stronger representation of body signals, namely (1) more accurate heartbeat-tracking ability, (2) stronger representation of the HEP, and (3) stronger experiential representation of the body (as measured with the SAQ) will be associated with more accurate duration reproductions.

## Results

### Behavioral findings

Two participants were excluded from further analysis because their data suggested that they were not paying attention to the task and reproduced and reacted randomly (more than 15% missing responses). Figure 1a represents the distribution of the mean reproduced durations for each interval. The general tendency towards under-reproduction increased with increasing interval duration (*M*(4s) = 3.95s, *SD*(4s) = 0.51; *M*(8s) = 7.41s, *SD*(8s) = 0.93; *M*(12s) = 9.71s, *SD* (12s) = 1.37). A repeated-measure ANOVA with the Greenhause-Geisser correction (F (1.19, 32.31) = 561.9, p < 0.001, η^2^= 0.95) showed that mean reproduced durations for 4s intervals were significantly shorter than 8s and 12s intervals (p < 0.001, p < 0.001, respectively), and reproductions of 8s were shorter than 12s intervals (p < 0.001).

**Figure 1.**
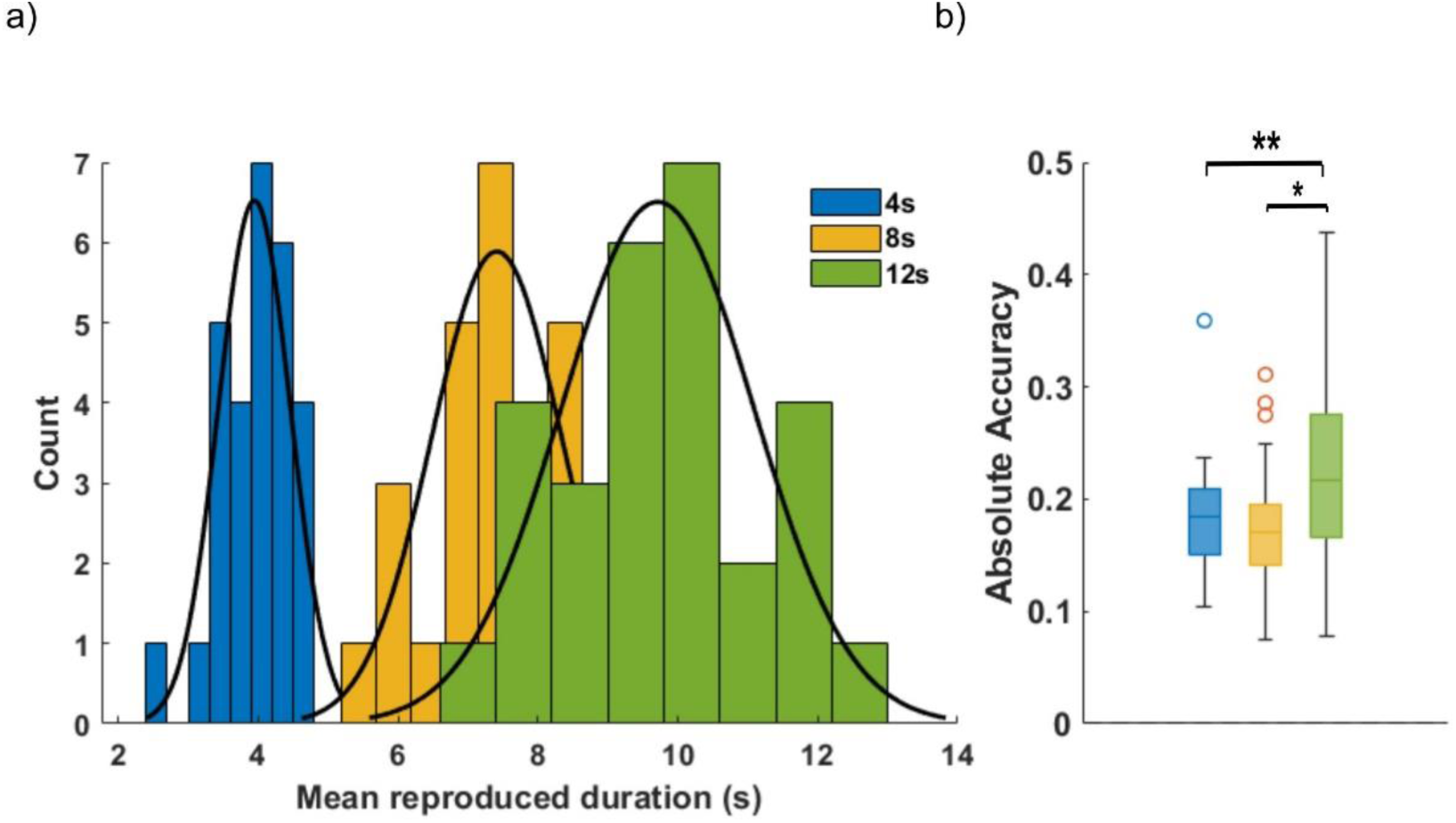
a) Histogram and density curves of the mean reproduced durations for different intervals. b) Boxplot illustrating the accuracy scores across different intervals. Each box represents the interquartile range (IQR) with the line inside indicating the median. The outliers are represented as individual points which are more than 1.5 * IQR away from the top or bottom of the box.

Regarding the absolute accuracy of the reproductions, there were significant differences between the different interval durations. Since the distribution of absolute accuracy was not normally distributed, analysis of variance was conducted using the Friedman Test, which showed a significant effect (χ² = 9.92, df = 2, *p* = 0.007), yielding a moderate effect size of Kendall’s W = 0.177. While the absolute accuracy did not differ significantly between 4s and 8s (p = 0.47), it was significantly higher (less accurate) in the 12s interval compared to the 4s (p = 0.013) and 8s intervals (p = 0.002) (Figure 1b). We observed a significant effect of interval on the precision of reproduced durations in terms of standard deviation (F (1.04, 28.08) = 1182.08, p < 0.001, η^2^= 0.97). The precision of reproduction declined significantly with interval length (all p < 0.001). The coefficient of variance of the reproduced durations also varied between intervals (F (2, 54) = 3.57, p = 0.035, η^2^= 0.11) with significantly lower coefficient of variance for the 8s intervals compared to the 4s ones (p = 0.031).

The mean accuracy of the secondary working-memory task during both tasks was relatively high (97.8% during the reaction time and 96.5% during the timing task), showing that participants did attend to the working-memory task, and this controlled for chronometric counting. The performance of the working-memory task did not correlate with the reproduced durations in all three intervals (p = 0.75, p = 0.42, and p = 0.84, respectively).

The accuracy of the heartbeat-counting task did not associate with the three reproduced durations (p = 0.48, p = 0.92, and p = 0.98). We observed positive correlations between the SAQ scores for interoceptive awareness and the reproduced durations. After correcting for three comparisons with the false-discover-rate (FDR) method (Benjamini & Hochberg, 1995), the SAQ score significantly correlated positively with the mean reproduced duration of the 4s interval (*r* = 0.40, *pfdr* = 0.031), 8s interval (*r* = 0.46, *pfdr* = 0.019), and 12s interval (*r* = 0.49, *pfdr* = 0.019). Negative correlations also emerged between the absolute accuracy values and the SAQ scores, but they were only significant for the 12s interval (*r* = -0.46, *pfdr* = 0.042). The larger the SAQ score, the smaller the deviation from the stimulus duration. This indicates that people with higher SAQ values were not only more accurate, but simultaneously showed stronger relative over-reproduction of the durations.

### Duration encoding versus reaction-time phases

Figure 2 presents the grand-average ERPs time-locked to the first sound in the reaction-time task and the encoding phase of the duration-reproduction task. The CNV is different for the encoding phase compared to the reaction-time task with more significant negativity for the timing task. The cluster-based permutation tests revealed that the observed CNV was significantly larger (more negative) within the 1000-4000ms time interval for 4s, 2100-5500ms for the 8s, and 3400-5300ms for the 12s duration (p < 0.05). The scalp topographies in these time intervals showed larger fronto-central negativity for the encoding phase compared to the reaction-time task. Interestingly, the significant time windows were around the offset of the shortest target duration (4s) when the brain predicts a likely offset.

**Figure 2.**
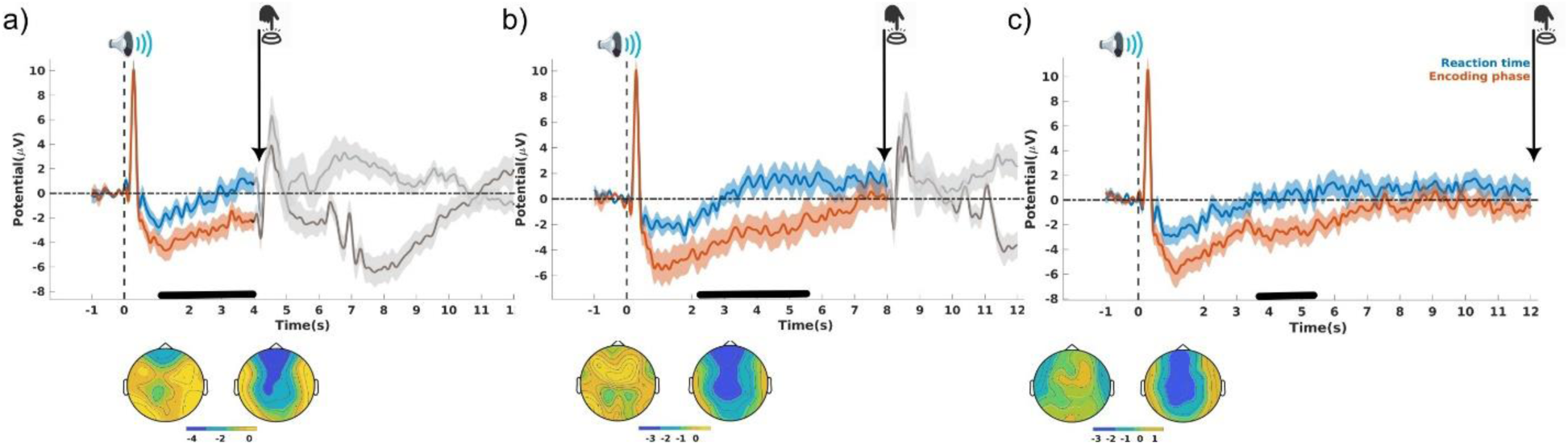
The upper plots (a-c) show the grand-average ERPs from the fronto-central scalp electrodes time-locked to the first sound in the reaction-time task (blue) and the encoding phase of the duration-reproduction task (orange) for the 4s (a), 8s (b), and 12s (c) intervals. The error shade represents the standard error, and the gray traces represent the interval portion beyond the target interval. Black traces in the time axes show the time interval in which the difference between two ERPs was significant according to the cluster-based permutation test. The lower plots (a-c) present the corresponding topography for the ERPs within these intervals (corresponding to the phases of the reaction-time task and duration encoding, respectively).

We explicitly extracted the CNV components (initial CNV or iCNV and late CNV or lCNV) to assess the modulation of CNV more precisely by both tasks. Repeated-measure ANOVA showed that the iCNV amplitude was significantly larger for the encoding phase of the duration reproduction than the identical stimulus-presentation phase in the reaction-time task (F (1,27) = 9.96, p = 0.004, η^2^ = 0.16) (see Figure 3a).

**Figure 3.**
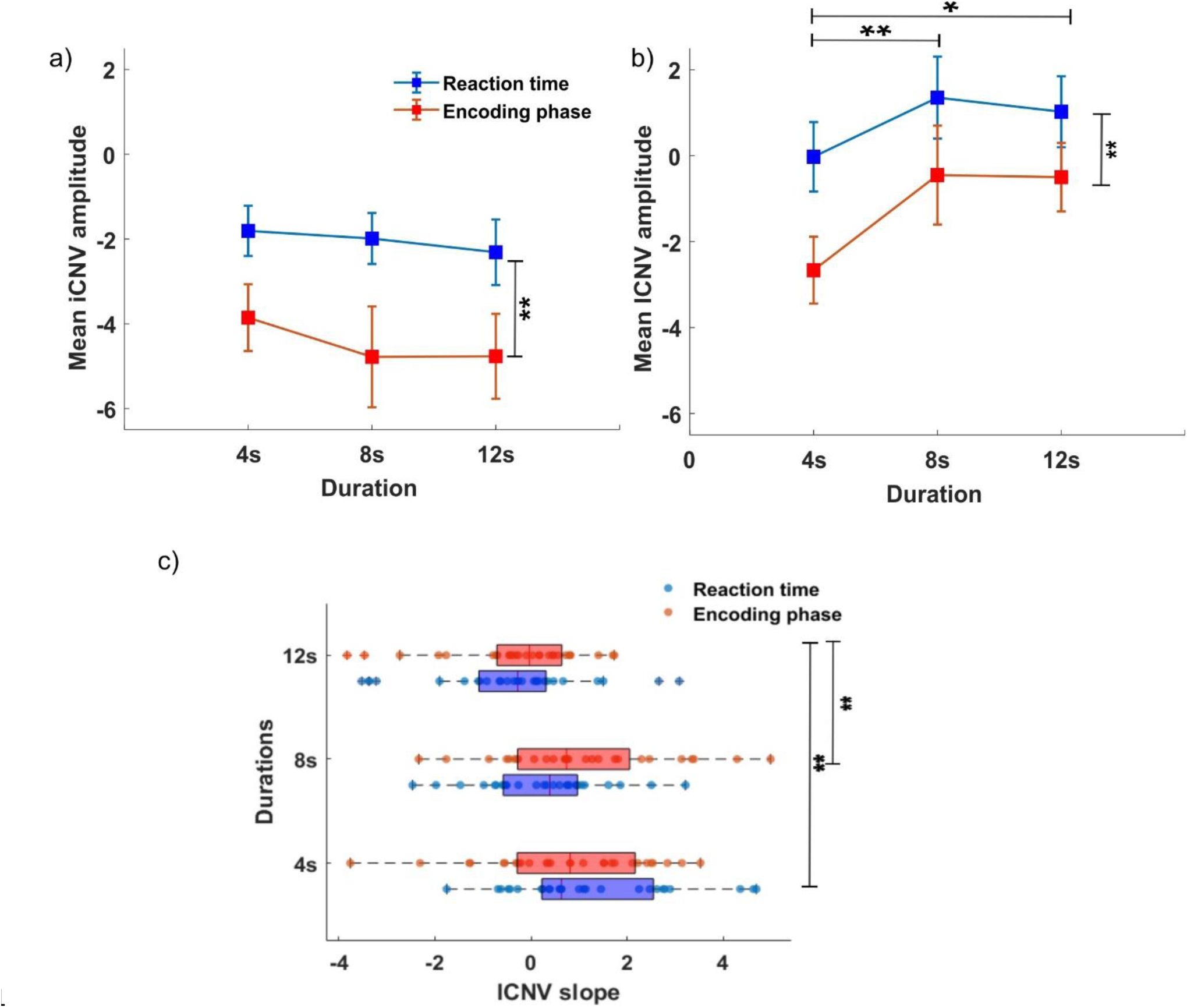
a) The average iCNV amplitude with corresponding standard errors for the reaction-time task and the encoding phase of the duration-reproduction task. b) The average lCNV amplitude with corresponding standard errors for both tasks. c) The average CNV slopes for both tasks and three intervals.

This indicates a neural-processing difference between these two tasks. After the onset of the first sound in the encoding phase, participants probably oriented their attention toward time and started storing temporal information in the working memory (to be able to reproduce the interval later) while waiting to press the button as soon as the sound ended (explicit timing). In the reaction-time task, they likely waited for the sound to finish and then pressed the button. No duration effects or interactions of duration and task were observed. We then assessed the lCNV amplitude by averaging the amplitudes of the CNV signal within the last 2 seconds of the interval. The main effects of task and duration were significant (F (1,27) = 10.21, p = 0.004, η2 = 0.089; F(2,54) = 5.01, p = 0.01, η2 = 0.058).

As shown in Figure 3b, the lCNV amplitude was significantly larger (more negative) for the encoding phase than for the reaction-time phase. Regardless of the task, the lCNV amplitude was more negative for the 4s intervals than for the 8s and 12s intervals (p = 0.01, and p = 0.02, respectively). Corresponding with the previous literature, larger lCNV amplitudes were associated with shorter durations [41,42]. As for the slope of the CNV signal (Figure 3c), we observed a significant main effect of duration (F (2,54) = 10.08, p < 0.001, η2 = 0.125). The slope of the CNV was significantly smaller for 12s intervals compared to the 4s (p < 0.001) and 8s (0.004) intervals.

### Duration reproduction versus reaction time

To compare the reproduction phase of the timing task with the reaction-time task, the ERPs were extracted in both tasks time-locked to the motor response (button press in the reaction-time task and button press in the reproduction phase of the timing task). The extracted grand-average ERPs are presented in Figure 4. Overall, the CNV signal before pressing the button during the duration-reproduction phase was larger (more negative) compared to the control task with significantly higher negative values for the 8s interval (cluster-based permutation test, p < 0.05). A clear tendency was also demonstrated for the 4s interval, but the difference was not significant.

**Figure 4.**
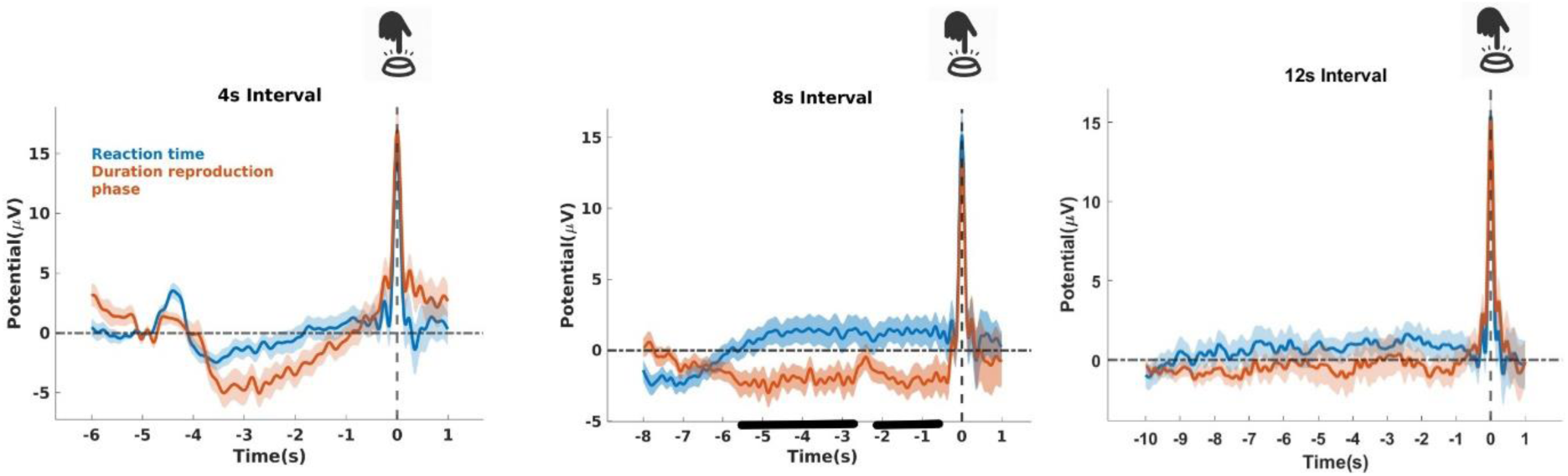
The grand-average fronto-central ERPs time-locked to the button press in both the duration-reproduction phase (red trace) and the reaction-time phase (blue trace).

We assessed the initial and late CNV amplitudes during the duration-reproduction phase. For this, the EEG signals were epoched time-locked to the reproduction cue (onset of the second sound) within 1s before to 12s afterward, and the iCNV, the lCNV, and slope of the CNV were calculated (Figure 5a). While the iCNV amplitude was extracted as an average CNV amplitude within 500-1500ms after the stimulus onset, the lCNV was identified as the average amplitude within the last 2s of the reproduction phase. The last 2s were also considered for the slope calculation. Considering the iCNV and lCNV amplitudes, the main effect of duration for both measures using a linear contrast was significant (F (2,54) = 2.90, p = 0.023, η2 = 0.097, F (1.58, 42.66) = 3.14, p = 0.036, η2 = 0.10). The following post-hoc tests revealed a marginally more negative iCNV and lCNV for 12s durations compared to 4s durations (p = 0.06, p = 0.09, respectively). For the CNV slope, the main effect of duration was significant (F (1.35, 36.49) = 8.42, p = 0.003, η2 = 0.23), highlighting significantly steeper slopes for the 4s interval compared to the 8s and 12s interval reproductions (p = 0.006 and p < 0.001, respectively). We assessed possible associations between the extracted CNV components and the reproduced durations. The reproduced durations showed a significant negative correlation with the mean lCNV amplitude (r = -0.46, p = 0.014) only for the 12s interval (Figure 5e). There was no correlation between the slope of the CNV and mean reproduced durations.

**Figure 5.**
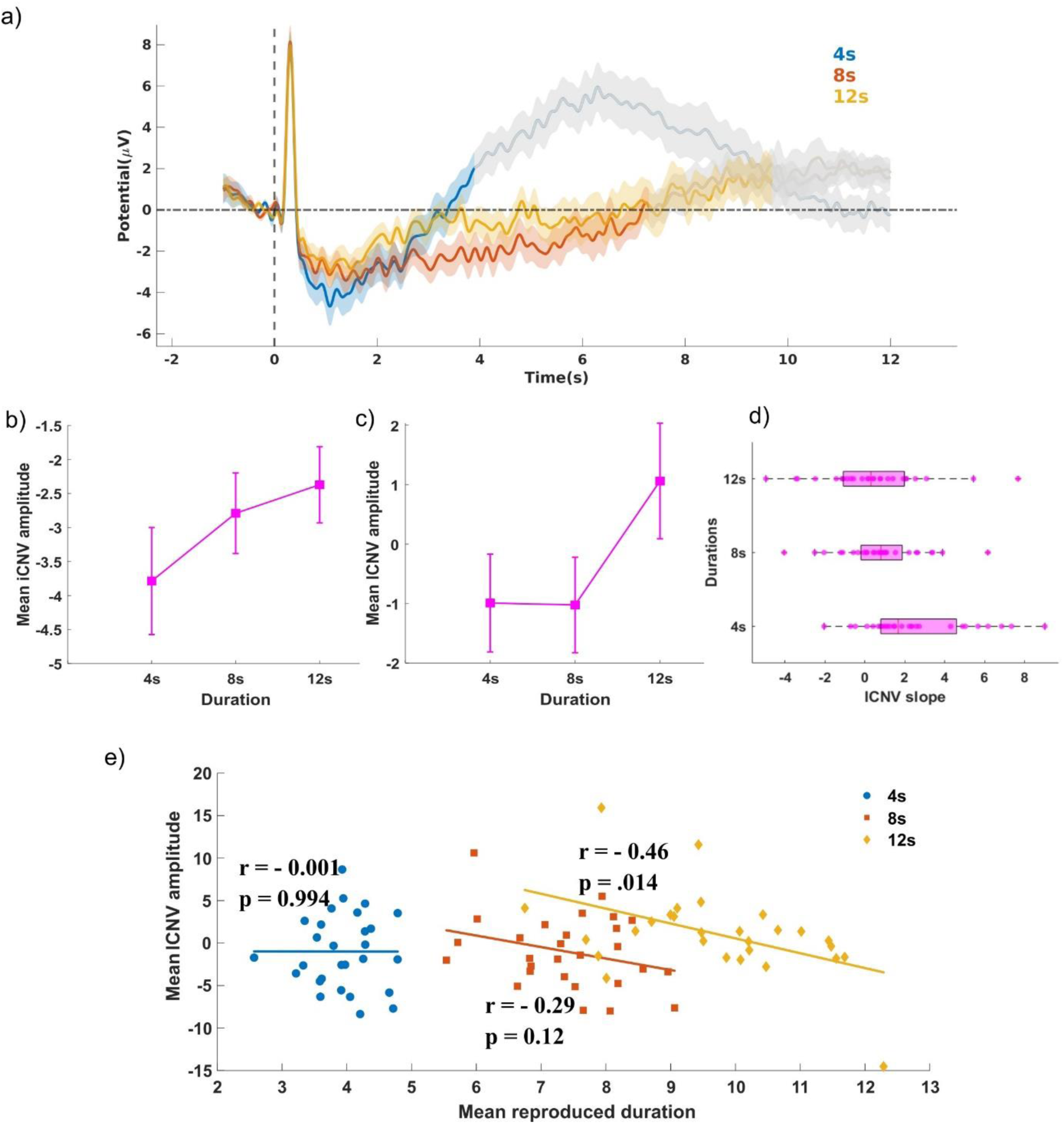
a) The grand-average fronto-central ERPs time-locked to the button press of the duration-reproduction phase. b) Mean iCNV amplitude with corresponding standard errors for different durations of duration reproduction. c) Mean lCNV amplitude with corresponding standard errors for different durations of the duration-reproduction phase. d) The average CNV slope for the three different durations of the duration-reproduction phase. e) The association between mean reproduced durations and the mean lCNV amplitudes during the reproduction phase.

### Modulation of the heartbeat-evoked potential during duration estimation

For each subject, the HEPs were averaged based on the time interval that occurred (HEP-4s, HEP-8s, and HEP-12s) for each task (the reaction-time task and the duration-encoding and duration-reproduction phase of the timing task). These grand-averaged HEPs were then compared across time intervals within each task using the cluster-based permutation test. The duration-encoding phase showed significantly different HEP values for the three durations within [130-270ms] and [470-520ms] after the R-peak (p < 0.05, cluster-based permutation test). The grand-average fronto-central HEPs for the encoding phase and significant time windows are presented in Figure 6. The HEP values around these time windows were significantly smaller for the 4s intervals than the 8s and 12s intervals. Given that no differences were observed for the reaction time or the reproduction phase of the duration-reproduction task, this highlights a duration encoding-specific function for the HEP amplitude within these time windows. The significant time windows correspond well with the latencies reported for the HEP modulation effects [43]. The average scalp topographies over the first- and second-time windows revealed less negative fronto-central HEP values for the 4s-encoding intervals compared to the other two intervals (see Figure 6). The average HEP in the first- and the second-time window are referred to as HEP1 and HEP2, respectively.

**Figure 6.**
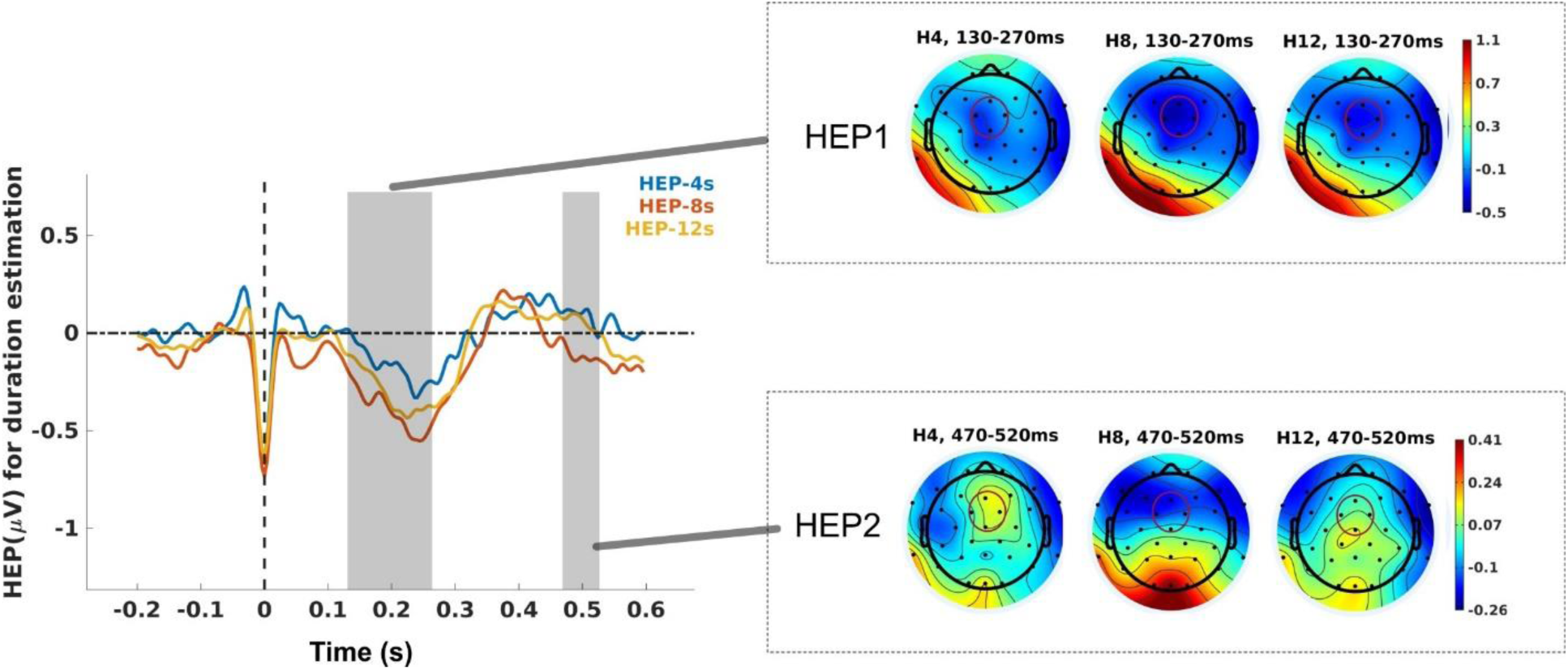
Grand-average HEPs correspond to the three target intervals in the encoding phase of the duration-reproduction task (left panel). Highlighted time windows show significantly different HEPs among the durations. The average topographies within the first-time (HEP1) and the second-time windows (HEP2) are presented in the right panel.

The HEP2 amplitude in the reproduction phase significantly correlated with the reproduced durations (r = - 0.43, p = 0.02) and with the absolute accuracy score (r = 0.46, p = 0.01) only for 8s intervals. This means that for the 8s interval, the higher the HEP2 amplitude during the reproduction phase, the shorter the durations reproduced by participants and the more inaccurate their responses.

### Gradual dynamics of the HEP components

#### Duration encoding vs. reaction time

Second-by-second dynamics of the HEP components (HEP1 and HEP2) for both tasks (the reaction-time task and encoding phase) separately for each time (4s, 8s, or 12s) are presented in Figure 7. We compared the HEP1 and HEP2 values between the two tasks (task effect) for each second (time effect) using the LME models. For the HEP1 values, a significant main effect of time (F(3,216) = 4.72, p = 0.008) was observed for the 4s intervals showing lower HEP1 values in the 2^nd^ and 3^rd^ seconds compared to the first second (p = 0.024, p < 0.001). Subsequent post-hoc tests showed that the HEP1 values decreased significantly from the first second up to the third second of the interval only for the encoding phase of the timing task (p = 0.027, p < 0.001).

**Figure 7.**
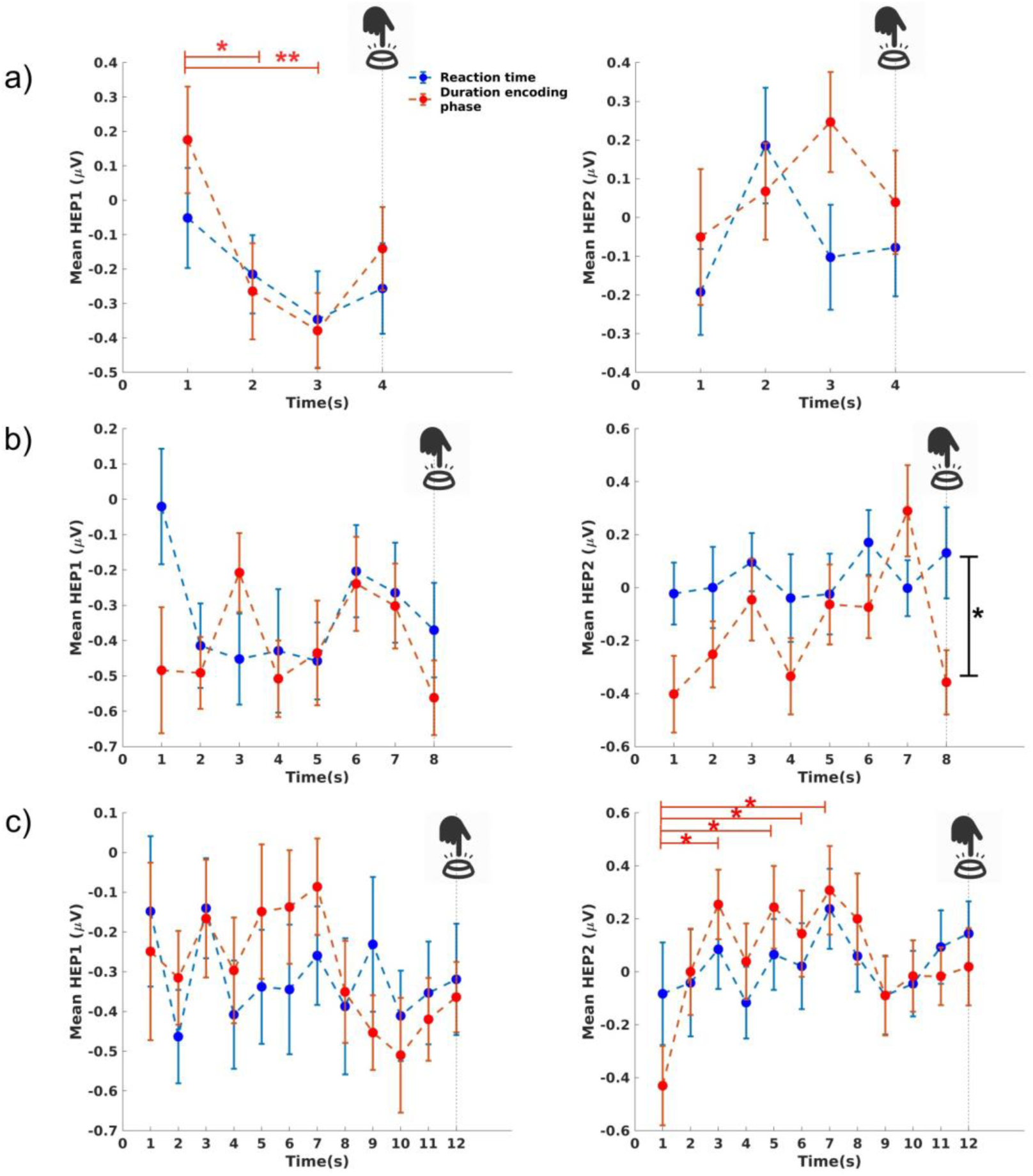
Second-by-second HEP1 (left column) and HEP2 (right column) amplitudes for a) 4s interval, b) 8s interval, and c) 12s interval during the reaction-time task and the duration-encoding phase. All significant differences are demonstrated with black and red lines. The red lines show that the corresponding difference was only significant in the encoding phase of the duration-reproduction task.

While the HEP1 values decreased second by second, the HEP2 values showed a rising trend for all three intervals. The LME model revealed a main task effect for the 8s interval (F(1,432) = 7.12, p = 0.007), reflecting more negative HEP2 amplitudes for the duration-encoding phase than the reaction time. A main effect of time was observed (F (11,648) = 1.90, p = 0.035) for the 12s intervals, showing higher HEP2 values for the 3^rd^, 5^th^, 6^th^, and 7^th^ seconds compared to the first second, but only for the encoding phase of the timing task (all p < 0.05). The slope of this gradual development for HEP1 and HEP2 showed no significant association with the reproduced durations. Similarly, the rate of change (difference between the first and last second HEP1 (HEP1diff) or HEP2 (HEP2diff) showed no correlation with the reproduced durations.

The cumulative HEP2 trend for each interval and each task was examined (Figure 8) to better understand the gradual change of the HEP2 component within each interval. Starting from the first second of the interval, an apparent ramp-like increase in the cumulative HEP2 amplitude could be observed during the 8s and 12s intervals for the encoding phase of the timing task, but not for the reaction-time task. We compared the cumulative HEP2 values between the two tasks separately for each interval using the LME models with task (reaction time or duration encoding) and time (first second up to last second of the interval) as fixed-effect factors. The LME model showed significant effects of the cumulative HEP2 for the 8s and 12s intervals. The cumulative HEP2 amplitude was significantly more negative for the encoding phase of the duration-reproduction task compared to the reaction-time task (F(1,420) = 52.54, p < 0.0001) for the whole 8s interval. No significant effect of time was observed for the 8s interval, indicating that the HEP2 amplitude does not differ as a function of time, although a trend is visible (Fig. 9b). Considering the cumulative HEP2 values within the 12s interval, the main effect of time was significant (F (11,648) = 3.38, p < 0.001). Post-hoc tests showed that the cumulative HEP2 values decreased/changed significantly from the third second up to the end of the interval for the encoding phase of the reproduction task (all p < 0.005).

**Figure 8.**
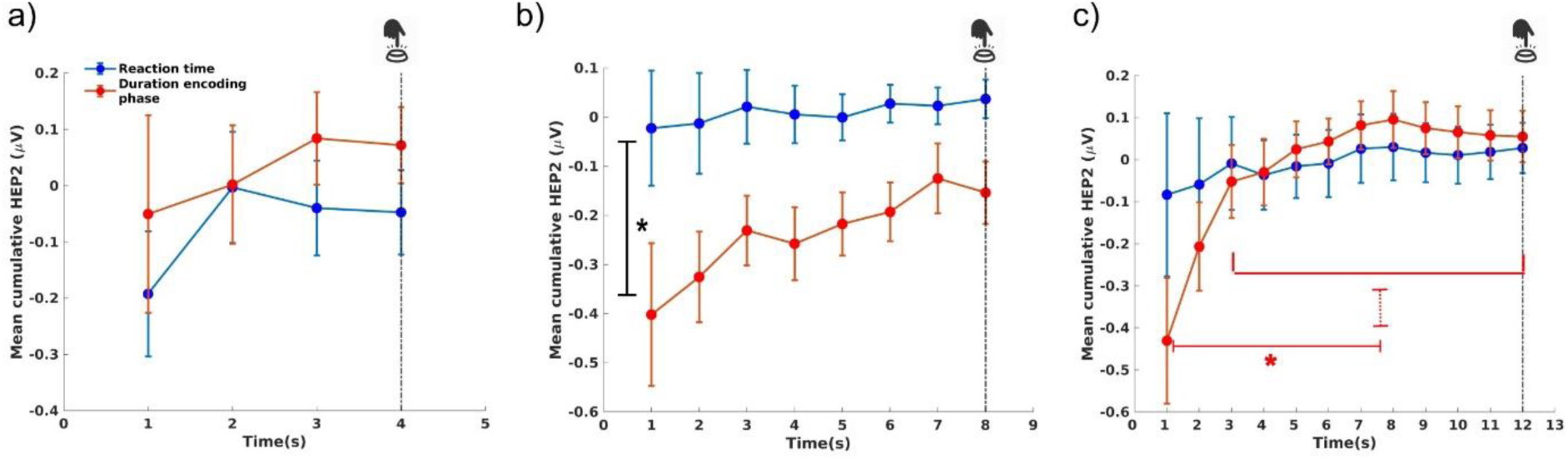
The cumulative HEP2 trend for the reaction time (blue line) and the encoding phase (red line). The solid lines show significant effects during the 8s and 12s intervals.

**Figure 9.**
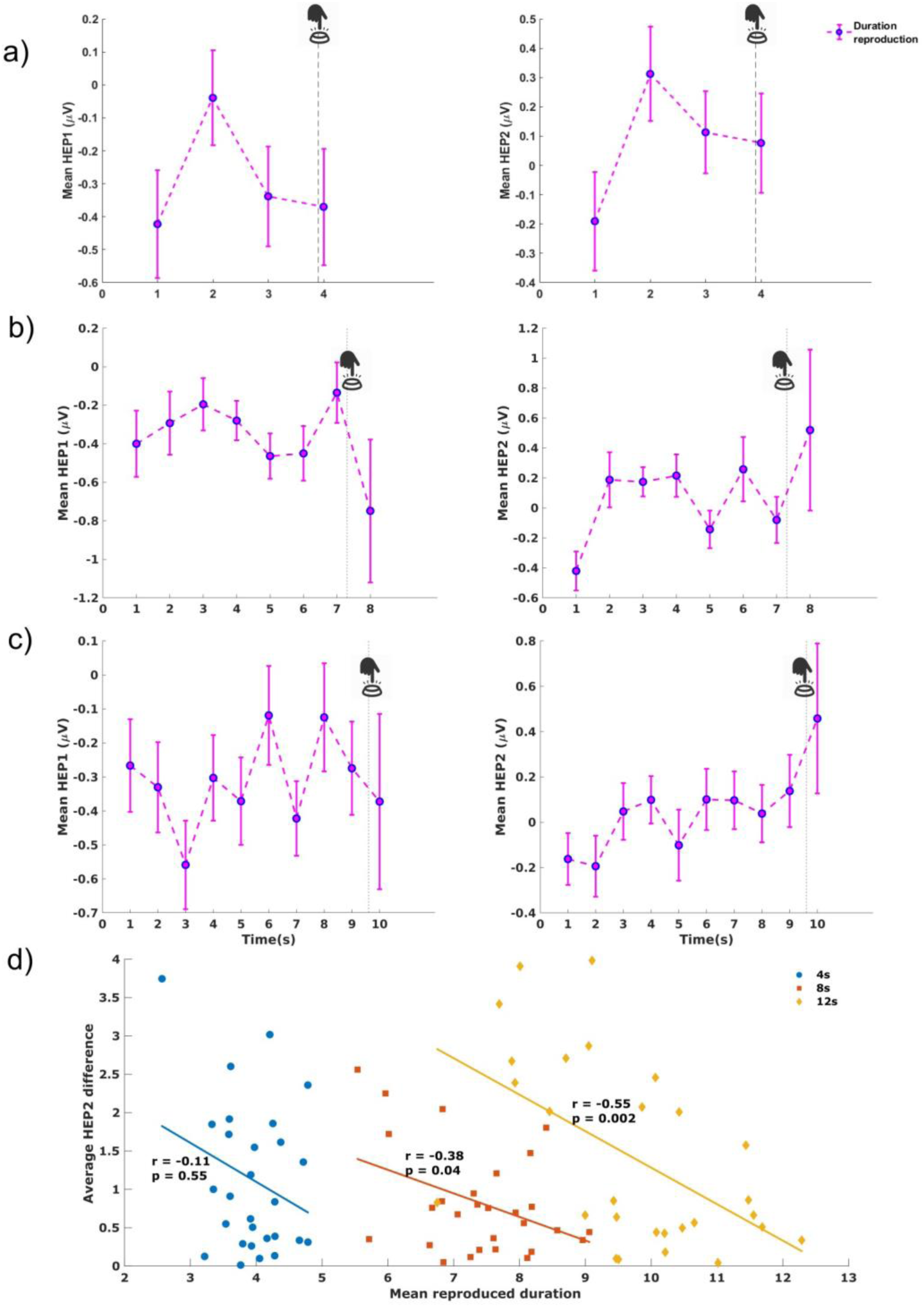
Second-by-second development of the HEP1 and HEP2 amplitudes for the duration-reproduction phase during the a) 4s, b) 8s, and c)12s intervals. d) The average change in the HEP2 amplitude between the first and last seconds during the reproduction phase correlated negatively with the reproduced duration in the 8s and 12s intervals.

#### Duration reproduction phase

We finally looked at the second-by-second development of the HEP components for the duration-reproduction phase. A similar gradual increase in the HEP2 was observed for all three intervals (Figure 9). The LME model did not show any significant main effect of time for each of the three intervals. Correlation analysis showed a significant association between the amount of change in HEP2 in the reproduction phase and reproduced durations for the 8s and 12s intervals (r = -0.38, p = 0.04; r = -0.55, p = 0.002).

The larger the HEP2 differences between the first and last seconds of the intervals (HEP2diff), the shorter the reproduced durations (Figure 9d).

### Neural indices predict duration reproduction

We performed a stepwise regression analysis to identify the significant neural indices that predict temporal reproductions in each target duration. We first fitted the model for each target interval with one independent variable (SAQ scores) and added other variables (lCNV amplitudes, CNV slope, HEP1diff, and HEP2diff) as possible predictors. The significance level for entry into the model was set at p < 0.05. Table 1 shows the final stepwise regression models for each interval-reproduction category along with the corresponding coefficients and associated statistics for the variables. The final model for the 4s-interval reproductions only included the SAQ scores and was statistically significant (F(26) = 5.21, p = 0.030), explaining 16.7% of the variance in the reproduced durations.

**Table 1.**
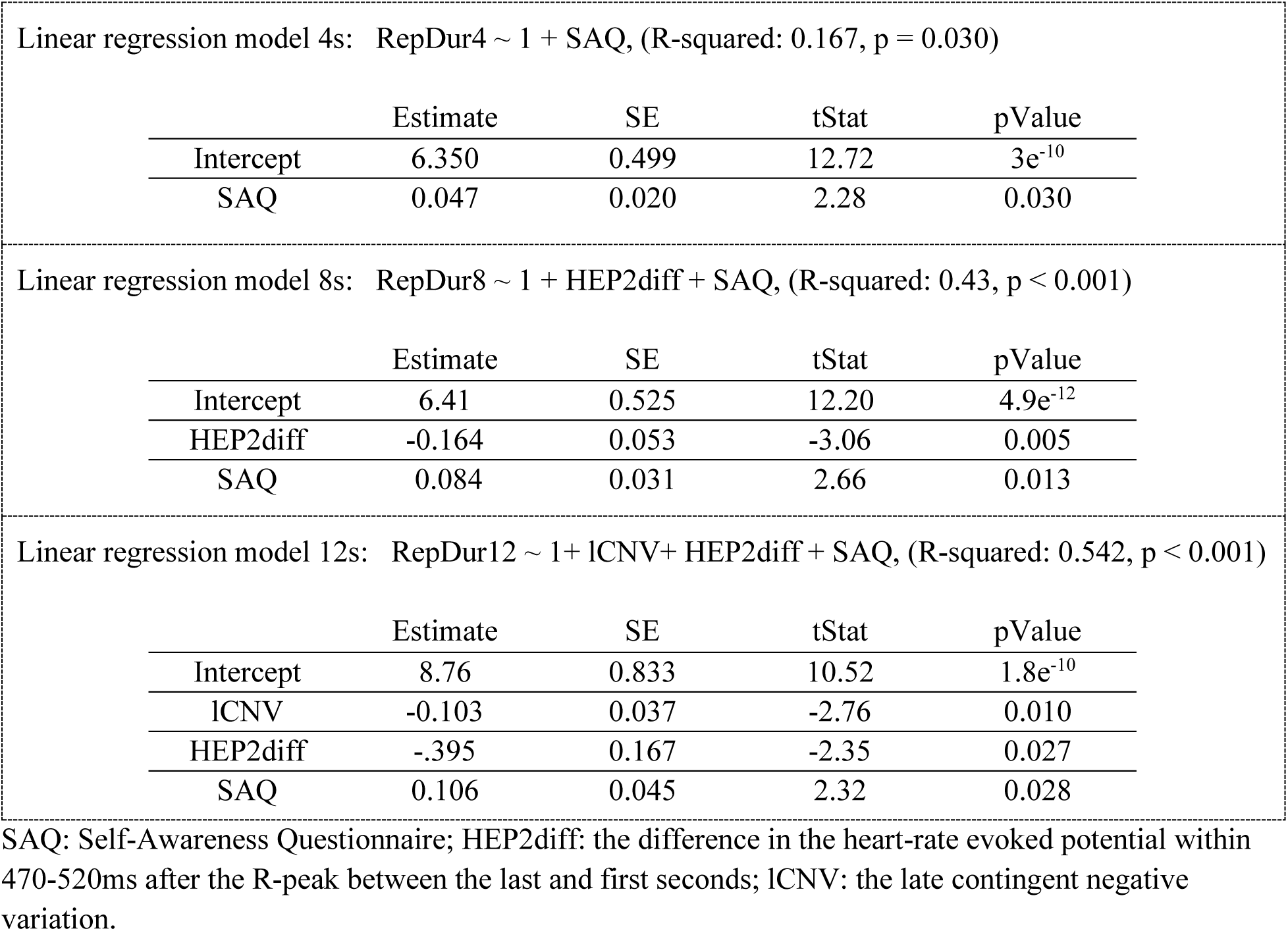
Results of the stepwise regression analysis.

The 8s-interval reproductions (the final model, which included the HEP2diff and SAQ scores as predictors), explained 43% of the variance in the reproduced durations (F(25) = 9.43, p < 0.001). Considering 12s intervals, the final model included the lCNV amplitude, HEP2diff, and SAQ scores as predictors and explained 54.2% of the variance in the reproduced durations (F(24) = 9.46, p < 0.001). The results of the stepwise regression analysis suggest that HEP2diff and SAQ scores were associated with the reproduced durations in the 8s and 12s intervals.

## Discussion

This study aimed to assess the association between time perception and interoceptive processes. To address this, we asked participants to perform a duration-reproduction task and a reaction-time task with 4s, 8s, and 12s intervals. The interoceptive ability was measured using the heartbeat counting task (HCT), the interoceptive Self-Assessment Questionnaire (SAQ), and the analysis of heartbeat-evoked potentials. Our behavioral findings confirmed previous studies (e.g., Eisler, 1976; Noulhiane et al., 2009; Wittmann et al., 2010) by showing that participants progressively under-reproduced intervals in relation to the target-interval length. A significant drop in reproduction accuracy was observed between the 4s and the 12s intervals and between the 8s and the 12s intervals. The longer the duration, the harder it was to retain its representation in working memory and reproduce it accurately.

The interoceptive awareness score (assessed by the SAQ) correlated positively with the mean reproduced durations for all intervals. Similar significant correlations were also observed for the absolute accuracy of reproductions of the 12s interval and the SAQ. This reflects that people with a lower SAQ score (less interoceptive awareness) under-reproduce the durations to a higher degree and are more inaccurate. Differently stated, the higher the score (a stronger awareness of bodily signals), the longer the duration reproductions and the more accurate the timing behavior. This association aligns well with the previous work showing that interoceptive awareness predicts timing accuracy in an irregular, but not in a regular, externally-cued condition [8]. As no regularly occurring external cues were present in this experiment (and no counting or rhythm-based strategy was used, as evidenced by the accurate performance of the secondary working-memory task), participants had to rely on interoception of body signals for timing intervals. However, this study failed to find any association between interoceptive accuracy derived from the HCT and the reproduced durations across all intervals, which stands in contrast to previous positive findings [8,17]. This discrepancy can partly be explained by different HCT paradigms that have been used. Teghil and colleagues applied an active motor-tracking, heartbeat-detection task in which participants were required to tap a key in synchrony with their heartbeats – a more objective task to study interoceptive accuracy (Garfinkel et al., 2022).

These outcomes relating to a partial association between interoception and subjective time are complemented by the results derived from the HEP analysis. The heartbeat-evoked potential (HEP) is an ERP component of the heartbeat that covaries with bodily self-consciousness [22,23]. Specifically, the HEP amplitude in the 200–600ms window after the R-peak is an index of cortical processing of afferent cardiac signals. We found a duration-specific modulation of this HEP signal only during the encoding phase of the duration-reproduction task. The HEP amplitude within interval borders [130-270ms] and [470-520ms] after the R-peak was smaller for the 4s interval than the 8s and 12s intervals. We named the average HEP within the first- and second-time windows HEP1 and HEP2, respectively. Given that no differences between intervals were observed in the reaction-time task, these results highlight a duration-specific function of the HEP signal. The HEP amplitude during these time windows increased with increasing time intervals in the encoding phase, which corresponds well with the results of the study by Richter & Ibáñez (2021). They demonstrated that the HEP amplitude was significantly higher within the borders of 182–222ms after the R-peak in those participants who overestimated duration compared to the under-estimator subgroup. We observed a negative correlation between the HEP2 value during the reproduction phase of the timing task and the reproduced durations for the 8s intervals. The larger the HEP2 amplitude, the shorter the reproduced interval.

The novelty of our study lies in our assessment of the gradual development of the HEP components in each task. The gradual development of the HEP2 amplitude presented in Figures 8 and 9 (cumulative as well as second-by-second development) resemble the previously reported ramp-like increases in cardiac periods [17] or the climbing activity in the insular cortex as detected in fMRI [14] during the encoding and the reproduction phases of the timing task. This ramp-like increase was significant for the duration-encoding phase of the timing task and not for the reaction-time task particularly for 12s intervals. While participants were trying to estimate the 12s interval, the HEP2 amplitude (especially the cumulative HEP2) gradually increased, showing a higher HEP2 after the third second up to the end of the interval compared to the first 3 seconds. These observations are consistent with the notion of the cortical accumulation of heartbeats as the neural mechanism of supra-second timing [13]. We also observed indications of a reset of the climbing HEP activity as a function of timing, which was most clearly visible in the 8s-encoding condition (see Fig. 9b). Climbing HEP amplitudes decreased just before the expected end of the stimulus, i.e., between 3 and 4 seconds, and then again between 7 and 8 seconds. No further climbing activity was observed for the encoding period between 8s and 12s, perhaps because the brain could predict the longest interval’s duration immediately once the sound did not stop at 8s. These preliminary findings and speculative interpretations should be studied further in follow-up studies.

Notably, the amount of HEP2 increases within the reproduction phase of the duration-reproduction task significantly correlated with the reproduced durations for the 8s and 12s intervals. The larger the increase in the HEP2 amplitude, the faster the subjective time and the greater the under-reproduction of duration. This is a striking correlation between the HEP2 as a measure of cortical sensitivity and behavioral timing.

Regarding the CNV amplitude, which has been suggested to represent an accumulation mechanism in the timing of duration [32,41,46,47], the timing task (the encoding phase and reproduction phase) led to significantly more negative CNV values (as tested by the cluster-based permutation test) compared to the control task. The average iCNV and lCNV components were significantly larger (more negative) for the encoding phase of the duration-reproduction task than for the reaction-time task. Both components (the iCNV and lCNV) have been linked to time perception, orienting attention, and early anticipation processes [41,42]. Given the link between these CNV components and attention modulation, we interpret these findings as a sign of stronger sustained attention towards time during the timing task. While no significant differences in the iCNV component were found for different durations, the lCNV amplitude covaried with the duration of the interval, reflecting the previously reported larger lCNV amplitudes for shorter durations [41,42]. A similar covariation was observed for the slope of the CNV, reflecting that shorter durations were associated with the larger CNV slope. The CNV slope was significantly higher for the encoding phase of the duration-reproduction task than for the reaction time, especially during 8s intervals.

Building on the aforementioned conceptual and empirical work, which the present results of our study complement, one can conclude that the passage of time is constituted through the existence of the bodily self across time as an enduring and embodied entity [12], an idea that was voiced by Merleau-Ponty (1945) in his phenomenological analysis: the physical self and subjective time are inseparable. The cognitive pacemaker-accumulator model with its abstract p̔ ulses’ can now be complemented with concrete entities: the neural signals from the body as subjectively experienced (as assessed with the Self-Awareness Questionnaire) and objectively measured by the heartbeat-evoked potential recorded 470 to 520ms after the R-peak. Accordingly, the two modulators of subjective time (attention and arousal) in the pacemaker-accumulator model can be understood as regulating the inflow of signals from the body, potentially as they accumulate over time in the insular cortex. A subjective expansion of duration is achieved through an increased awareness of bodily states or an increase in the amplitude of the HEP, either through more attention to the bodily self or through increased physical arousal.

The present study has some limitations that should be acknowledged. First, we restricted trial numbers to 126 (42 per target duration) for practicality, which potentially limited the statistical power of our analysis, particularly for the analysis of HEP dynamics across time. Next, we employed a secondary working-memory task to prevent the confounding effects of using a counting or a rhythm-based strategy for time estimation. Adding an additional, ecologically valid task without this constraint might have been beneficial. Finally, the small number of EEG channels (32 channels) recorded prevented performing source localization, which would have been very useful for relating the findings with previous fMRI studies that used the same paradigm. Future studies should use a higher-resolution EEG recording to produce a more accurate source localization. This work explored the function between CNV, HEP, and temporal reproductions within the supra-second range (4s, 8s, and 12s). Future studies should consider a broader range of intervals to ascertain the HEP’s relevance across different time scales. Longer intervals may invoke distinct cognitive strategies beyond the HEP (e.g., memory load) or slower interoceptive signals, such as the breathing rate.

Just how and where in the brain event duration is processed remains unanswered. Timing mechanisms in the brain are of the essence for an organism to represent environmental temporal regularities and the temporal metrics of events [48]. Based on conceptual considerations from functional neuroanatomy and from experimental psychophysiology, we suggest that interoceptive (bodily) states create the experience of time. Within this theoretical framework of the embodiment of time, we presented accumulating evidence of how interoception and associated brain networks process duration. We postulate that the ongoing creation of an embodied self over time by ascending neural signals from the body – forming the material m̔ could function as a measure of time.

## Materials and Methods

### Participants

We tested 30 healthy participants (16 males, 14 females) aged 18 to 36 years (mean age: 22.9, SD: 4.2 years) recruited via an online advertisement, flyers, and word-of-mouth. Before participants came to the laboratory, a brief telephone screening was administered to check the inclusion and exclusion criteria (e.g., they were asked about psychological or neurological disorders). Participants were appropriately remunerated for their time and participation in the study. The study was approved by the local ethics committee of the Institute for Frontier Areas of Psychology and Mental Health (IGPP_2021_05).

### Experimental design

The experimental session took approximately three hours. After being welcomed by the experimenter, participants were informed about the purpose of the study and the experimental procedure and provided informed consent for their participation. They then filled out the SAQ. The schematic illustration of the experimental session and tasks used is presented in Figure 10. Following a 5-minute baseline recording with their eyes open, participants performed a heartbeat-counting task and subsequently a duration-reproduction or a reaction-time task in a counter-balanced order (Figure 10a). The latter two tasks were designed in three blocks, respectively. During each pause between the blocks and between the tasks, participants were allowed to stand up, walk around, drink water, or go to the toilet if they wanted to. All tasks were programmed and implemented with the experimental software PsychoPy (Version 2021.1.4, Peirce et al., 2019).

**Figure 10.**
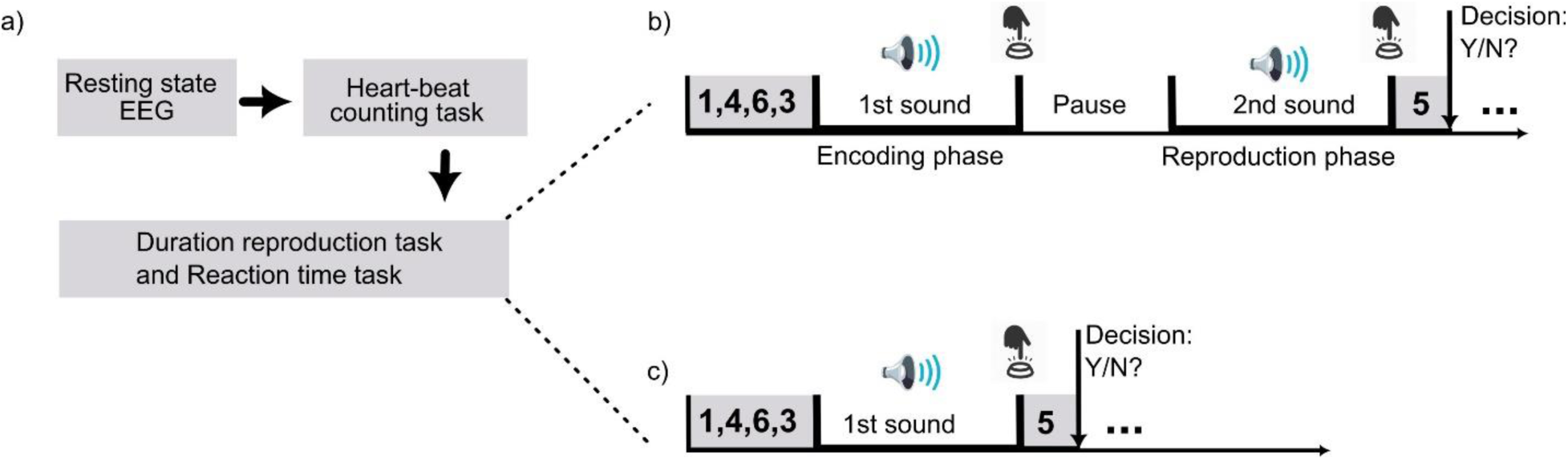
a) Experimental design of the study. The schematic illustration of the (b) duration-reproduction task and (c) reaction-time task. For details, see the methods section.

### Duration-reproduction task

The schematic illustration of the duration-reproduction task is presented in Figure 10b. Participants were first presented with an auditory stimulus (encoding phase; pink noise) lasting for a specific target duration (4, 8, and 12 seconds), which they then had to reproduce by pressing a button when they thought that a second comparison stimulus (reproduction phase; brown noise) was as long as the first one (Figure 10b). We used two different noises during the task to provide an additional cue indicating which phase of the task participants were in. There was a pause between the two stimuli with a jittered time interval between 2 and 3 seconds. Participants were also asked to react as quickly as possible at the end of the encoding phase by pressing a key. This reaction-time task had already been applied in the fMRI studies [14,15] and served to compare the encoding and the reproduction phases with the control reaction-time task (see below), respectively, when analyzing the EEG data in the present study. The auditory stimuli were created with the software Audacity (version 3.0.0, 2021). The pink and brown noises were chosen because they did not become unpleasant over time compared to pure sinus tones, which can become annoying. Different sounds were used during encoding and reproduction to help participants keep track of the trial structure; it can otherwise become confusing after prolonged testing.

Participants were instructed not to count or use any other rhythm-based strategy while exposed to the stimuli because chronometric counting yields more accurate and precise duration estimates and is governed by different brain processes [50,51]. Each trial included a secondary working-memory task to further prevent any counting. At the beginning of each trial, four different numbers from 1 to 8 which had to be memorized were shown on the screen for three seconds. At the end of each trial, one number was presented for three seconds, and the participant was then asked whether this number was one of the four digits shown at the beginning. Inter-trial intervals were set with a jittered duration between 1 and 2 seconds. A total of 126 trials were randomly presented with 42 trials allocated for each target duration.

### Control reaction-time task

During this task, auditory stimuli (pink noise) with 4-, 8-, or 12-second durations were presented, and participants were instructed to press a button as fast as possible when the stimulus ended (Figure 10c). The same secondary working-memory task explained in the duration-reproduction task was included in each trial. This made the control task comparable in timing-irrelevant cognitive load to the encoding phase. The 126 trials were presented in random order.

### Heartbeat-counting task (HCT)

Participants’ interoceptive accuracy, i.e., their ability to correctly perceive body signals, was assessed by a heartbeat-counting task [26]. Participants were instructed to count their heartbeats during given time intervals. They were requested not to monitor their pulse (e.g., by feeling their radial artery on the wrist), but by instinctively feeling their heartbeats. A beep indicated the start and the end of the interval while participants had to count their heartbeats. At the end of each time window, participants typed their heartbeat counts on a keyboard and indicated (with a mouse click on a visual slider scale) how confident they were, ranging between very unsure’ to very sure’. There were three training trials with randomly selected durations (25s, 30s, and 35s) and six experimental trials with durations of 25s, 30s, 35s, 40s, 45s, and 50s in randomized order. The objective heartbeat counts were recorded by electrocardiogram (ECG), which was part of the EEG acquisition setup.

### Questionnaires

The subjective awareness of body states was assessed with the German version (SAQ-17; Kübel & Wittmann, 2020) of the Self-Awareness Questionnaire (SAQ, Longarzo et al., 2015). Participants score how often they generally feel specific body sensations with answer categories ranging between 0 (‘never’) and 4 (̔always’) on 17 Likert-scale items.

### Signal recording

Continuous EEG signals were recorded using a 32-channel electrocap (ActiCHamp, BrainVision) with active electrodes positioned according to the extended 10-10 international system (‟American Electroencephalographic Society Guidelines for Standard Electrode Position Nomenclature,” 1991). All electrodes were referenced to the Fz electrode with the ground electrode placed on the forehead. Electrode impedances were kept below 10 kΩ. EEG signals were digitized with a 1000-Hz sampling rate and band-pass filtered within the 0.01–120-Hz range. One ECG signal was acquired using three Ag/AgCl electrodes, which were positioned according to the Lead II Einthoven configuration: two electrodes were placed on the right clavicle, and the left hip/abdomen (active electrodes), and one electrode was placed on the left clavicle (ground electrode). ECG signals were co-registered with the EEG via the amplifier’s auxiliary input.

### Data processing

The data processing was implemented with the Matlab software (R2022b, MathWorks) using custom-written scripts and the EEGLAB toolbox functions. After down-sampling the recorded EEG signals to 250 Hz and applying an FIR filter within the 0.05-70-Hz range, line noise was adaptively estimated and removed using a set of Sleppian filters or multi-tappers. After this initial filtering step, large non-stationary artifacts were identified and corrected using the Artifact Subspace Reconstruction (ASR) approach [54]. The cleaned EEG signals were then re-referenced to the average reference and decomposed into independent components (ICs) using the Adaptive Mixture Independent Component Analysis (AMICA; Palmer et al., 2006, 2011). Since the low-frequency EEG can affect ICA decomposition results [57], we calculated the ICs using 1-70-Hz band-pass filtered data and applied it to 0.05-70-Hz band-pass filtered data. The identified ICs were then automatically classified and labeled using the ICLabel plugin. This machine-learning approach has been trained to classify the ICs based on several characteristics, such as spectral properties and brain topography [58]. Non-brain-related ICs were selected for exclusion based on their spectra, scalp maps, and time courses. This procedure ensures cleaning of the EEG signals from eye blinks, muscle noise, heart artifacts, and other contamination. Epoch extraction was carried out separately based on the component of interest.

We re-referenced the cleaned EEG signals to the average of linked mastoids for the CNV signal analysis. To compare the reaction time and the encoding phase of duration-reproduction tasks, the EEG signals of both tasks were epoched around the onset of the encoding sound from 1s before to 12s after the onset. The average ERPs over the fronto-central scalp electrodes (Fz, FC1, Cz, FC2, F1, C1, C2, F2, and FCz) were then considered for the CNV signal comparison (for reference, see Kononowicz et al., 2015; Robinson & Wiener, 2021). We also contrasted CNV signals between the reproduction phase of the duration-reproduction and the reaction-time tasks. The fronto-central EEG signals of both tasks were first epoched, time-locked to the onset of the sound (encoding sound in the reaction-time task and reproduction sound in the time-reproduction task) with 3s before to 15s after the sound. The extracted segments were baseline corrected considering a 500ms pre-sound time window. From the obtained segments, final epochs were extracted time-locked to motor responses in both tasks (motor response to the button press) with 6s (for 4s intervals), 8s (for 8s intervals), and 10s (for 12s intervals) before the motor response to 1s after that. These intervals were chosen based on the mean reproduced time of these target durations.

To assess the initial CNV (iCNV) component, we calculated the average amplitude of the extracted fronto-central ERPs within 500-1500ms after the stimulus [41,59]. The late CNV (lCNV) component was considered as the average amplitude of the fronto-central ERP within the last 2s of each interval. The CNV slope as the slope of the fitted line to the ERP in this time window was also calculated.

For the HEP analysis, we first filtered the preprocessed signals using a high-pass filter with a cut-off frequency of 1Hz. After detecting the exact location of R-peaks in the ECG signal using discrete wavelet transform analysis with the “sym4” wavelets [60], their location was integrated into the EEG signals. Data epochs were then generated time-locked from -200ms before to 600ms after the detected R-peaks. Baseline correction was performed from –100 to –50ms. We focused on the average HEP signal over a fronto-central cluster (FC1, Cz, FC2, Fz), as reported in the previous literature [23,43,60]. Since we aimed to assess the association between the interval length and the HEP signal, we categorized each HEP signal based on the corresponding interval when it occurred (HEP-4s, HEP-8s, and HEP-12s). The HEPs were then averaged for each duration interval and each task. This procedure yielded three grand-averaged HEPs for each task (the reaction-time task, the encoding, and the reproduction phase of the duration-reproduction task). The average HEP amplitudes within time windows with significant differences are referred to as HEP1 and HEP2, respectively. Apart from the average HEP1 and HEP2, we assessed the gradual development of these components over time by calculating the rate of change across the whole target duration (HEP1diff and HEP2diff).

### Statistics

The dependent behavioral variables in the duration-reproduction task were reproduced durations, absolute accuracy, and precision (variance) scores derived from the reproduced durations. Absolute accuracy was defined as an absolute deviation of the reproduced duration from the presented stimulus duration divided by the presented duration. The standard deviation and coefficient of variance of the reproduced duration for each interval measured the participants’ precision of estimates. The following statistics were carried out with the statistical software Jasp (Version 0.17.2.1, 2023) and with Matlab (R2022b, MathWorks):

(1) One-way repeated-measures ANOVAs for within-subject differences or Friedman tests (for non-normality of the distribution) were calculated for behavioral comparisons across three target durations. (2) Differences in the fronto-central CNV or HEP signals between tasks or durations were tested using the cluster-based permutation t-test [61] implemented in the Fieldtrip toolbox [62]. With this procedure, samples with a *t*-statistic higher than a threshold (p < 0.05, two-tailed) are clustered in the same set based on temporal adjacency. The sum of the *t*-values within a cluster was assigned as a cluster-level statistic, and the null hypothesis was evaluated using the maximum of these cluster-level statistics. The two-tailed Monte-Carlo p-value to reject the null hypothesis corresponds to the proportion of cluster-based randomizations that resulted in a larger test statistic than the observed original cluster-level statistic. (3) We used a 2×3 repeated-measure ANOVA test with the Greenhouse-Geisser correction for degrees of freedom to compare the mean CNV and the mean HEP components considering two tasks (the reaction-time task and the encoding phase of the timing task) and three durations (4s, 8s, and 12s) as within-subject factors. (3) Correlations were calculated with Pearson’s correlation coefficients. Whenever we found a non-normality of the distribution in one of the two variables, the Spearman correlation coefficient (ρ) was used. (4) The gradual development of the HEP components was evaluated separately for each interval using the linear mixed-effect-models (LME) statistic. We considered the task (the reaction-time task and the encoding phase of the timing task) and time (the first second up to the last second of the interval) as fixed-effect factors with each participant as a random intercept. F-test results were obtained using ANOVA on the fitted model, and whenever the ANOVA results were significant, the pair-wise comparisons were calculated. (6) A final stepwise regression analysis was performed in Matlab using the six independent variables (iCNV and lCNV amplitudes, CNV slope, HEP1diff, HEP2diff, and SAQ) to identify the key factors influencing the reproduced durations. The significance level for entry into the model was set at p < 0.05.

## Acknowledgments

This study was funded by the EU, Horizon 2020 Framework Program, FET Proactive (VIRTUALTIMES consortium, grant agreement Id: 824128 to Marc Wittmann). VIRTUALTIMES — Exploring and modifying the sense of time in virtual environments — includes the following groups with the principal investigators Kai Vogeley (Cologne), Marc Wittmann (Freiburg), Anne Giersch (Strasbourg), Marc Erich Latoschik, Jean-Luc Lugrin (Würzburg), Giulio Jacucci, Niklas Ravaja (Helsinki), Xavier Palomer, and Xavier Oromi (Barcelona).

## Authorship contribution statement

SK: conceptualization, investigation, data curation, formal analysis, writing of the original draft, and visualization, DL: data curation, formal analysis, writing, review, & editing, MÇ and FA: data curation, writing, review, & editing, VN: conceptualization, writing, review, & editing, MW: conceptualization, funding acquisition, supervision, writing, review, & editing

## Data and code availability

The complete data supporting this study’s findings are available for researchers from the first author upon reasonable request. The example dataset and the code used for the analysis will be made available on the Open Science Framework website (https://osf.io/hj35a/?view_only=96d33ab8a9db4be0b60a271ca90a8f2c).

